# Unlocking the Reversal Potential of Solid Supported Membrane Electrophysiology to Determine Transport Stoichiometry

**DOI:** 10.1101/2020.05.07.082438

**Authors:** Nathan E. Thomas, Katherine A. Henzler-Wildman

**Affiliations:** Department of Biochemistry, University of Wisconsin-Madison

## Abstract

Transport stoichiometry provides insight into the mechanism and function of ion-coupled transporters, but measuring transport stoichiometry is time-consuming and technically difficult. With the increasing evidence that many ion-coupled transporters employ multiple transport stoichiometries under different conditions, improved methods to determine transport stoichiometry are required to accurately characterize transporter activity. Reversal potential was previously shown to be a reliable, general method for determining the transport stoichiometry of ion-coupled transporters (Fitzgerald & Mindell, 2017). Here, we develop a new technique for measuring transport stoichiometry with greatly improved throughput using solid supported membrane electrophysiology (SSME). Using this technique, we are able to verify the recent report of a fixed 2:1 stoichiometry for the proton:guanidinium antiporter Gdx. Our SSME method requires only small amounts of transporter and provides a fast, easy, general method for measuring transport stoichiometry, which will facilitate future mechanistic and functional studies of ion-coupled transporters.

## Introduction

Ion-coupled transporters utilize energy stored in electrochemical gradients to drive uphill substrate transport across cellular membranes. These transporters are essential to numerous physiological processes, from nutrient uptake to neural signaling, and are common drug targets, but successful therapeutic design is limited by our understanding of these integral membrane proteins’ structures, functions, and mechanisms (Garibsingh and Schlessinger, 2019; Lin et al., 2015). Transport stoichiometry – the number of ions and substrates moved per transport cycle – is a function of the transporter’s mechanism and is a crucial determinant of the transporter’s function, including the direction of driven transport, the energy spent per substrate transported, and the maximum substrate gradient that can be maintained at equilibrium. Generally, ion-coupled transporters have been assumed to operate according to tightly coupled mechanisms with a single, set transport stoichiometry. This understanding has led to the traditional classification of transporters as either antiporters, symporters or uniporters. However, there is growing evidence that many ion-coupled transporters operate through complex mechanisms that violate the assumption of stoichiometric transport, engaging in behavior that cannot be cleanly classified as antiport, symport, or uniport. (Bazzone et al., 2017b; Burtscher et al., 2019; Coady et al., 1996; Hussey et al., 2020; Kang and Hilgemann, 2004; Sacher et al., 2001). In light of these findings, there is a pressing need for detailed mechanistic investigations of a greater variety of transporters.

The structural revolution has revealed specific substrate binding sites on transporters, and together with biochemical assays, this has significantly advanced our understanding of transporter:substrate binding stoichiometry, (Guskov et al., 2016; Li et al., 2015; Piscitelli et al., 2010; Schwartz et al., 2005). However, it is important to distinguish binding stoichiometry from transport stoichiometry (Grabe et al., 2020). At times, ions or substrates can bind as allosteric effectors without being transported (Quick et al., 2018). This is especially difficult to untangle in the case of proton-coupled transporters with multiple protonatable side chains (Bozzi et al., 2019; Parker et al., 2014). Furthermore, establishing that substrate bound at a specific site can be transported is insufficient to establish that its transport is obligatory to the mechanism (Bazzone et al., 2017b; Robinson et al., 2017). Thus, measuring a transporter’s coupling stoichiometry not only sheds light on its biological function, but also provides crucial information about the transport mechanism.

The only way to reliably determine transport stoichiometry is through transport assays. Reversal potential assays have emerged as the method of choice for electrogenic transporters, as they are applicable to a variety of systems and allow for model-independent determination of the transport stoichiometry (Fitzgerald et al., 2017; Nguitragool and Miller, 2006). In brief, these assays follow transport as a function of membrane voltage for a set initial substrate and/or ion gradient. The reversal potential, the membrane voltage that results in no net transport, can be used to determine the transport stoichiometry. In theory, this method can be applied to any electrogenic transporter, but there are several important limitations. First, transport must be able to be monitored. Several eukaryotic transporters are amenable to patch clamp electrophysiology (Ravera et al., 2007; Shao et al., 2014; Zerangue and Kavanaugh, 1996), but this technique is generally unsuitable for low-flux transporters or for transporters that cannot be highly expressed in eukaryotic cells, including many structurally characterized prokaryotic transporters. Thus, we will focus on assays of purified protein in proteoliposomes, as these assays allow for characterization of the widest range of transporters. Ideally, liposomal transport can be followed through a real-time probe such as a fluorophore, but fluorophores are not available for every coupling ion. Radioactive transport assays may be used instead (Fitzgerald et al., 2017) but do not offer real-time monitoring, significantly increasing the time and effort required to collect data for the multiple time points and conditions needed to determine transport stoichiometry. A second issue with traditional liposomal assays is that liposome contents cannot easily be changed after reconstitution. This can make it difficult to be certain of the internal ion and substrate concentrations and increases the difficulty of screening a wide range of assay conditions. A third issue is that liposomal assays can require a large amount of sample, depending on the sensitivity of the detection method and the number of conditions that must be tested. Finally, for many of the reasons listed above, traditional liposomal transport assays are low throughput and require significant time and effort, even if the material costs can be kept to a minimum. As a consequence, transport stoichiometry been measured for only a small fraction of structurally characterized transporters (Fitzgerald et al., 2017; Groeneveld and Slotboom, 2010). A method for routine measurements of transport stoichiometry would greatly facilitate functional and mechanistic studies of ion-coupled transporters.

Here we present a new method for measuring coupling stoichiometry by adapting reversal potential assays to solid-supported membrane-based electrophysiology (SSME) (Bazzone et al., 2017a). SSME carries several advantages over traditional liposomal transport assays. By directly measuring transported charge, SSME allows real-time monitoring of electrogenic transport without the need for fluorophores or radiolabeled substrates. In addition, its sensitivity requires only picomole amounts of protein to achieve sufficient signal. Finally, dozens of conditions can be tested on a single sensor without the need for separate reconstitutions, greatly increasing the throughput of the assay. We demonstrate the utility of this method using *E. coli* Gdx, whose 2:1 proton:guanidinium antiport stoichiometry was recently established using traditional reversal potential measurements performed by another lab (Kermani et al., 2018). With our SSME assay, we confirmed this result, using less than 2 nmol total protein to acquire 200 transport assay measurements in under three days. In addition, we demonstrate that it is possible to change the internal ion and substrate concentrations using the SSME setup, enhancing the number of experimental conditions that can be tested on a single protein sample. This assay is fast, easy, and accurate, requires a minimal amount of sample, and is potentially applicable to any electrogenic transporter regardless of coupling ion.

## Results

Reversal potential assays rely on the ability of coupled transporters to perform reversible work. That is, transport of ions down large (electro)chemical gradients can drive transport of substrates up concentration gradients, and conversely, transport of substrates down large gradients can drive transport of ions up (electro)chemical gradients. When the electrochemical gradients are balanced, no net transport occurs, allowing for calculation of the transport stoichiometry (see *Materials and Methods* for equations and derivations). Our assay simply adapts this principle to SSME.

In an SSME experiment, proteoliposomes are adsorbed onto a membrane-coated gold electrode, creating the sensor. To initiate an experiment, buffer is run over the sensor in three stages. First, a “non-activating” buffer containing the same solution as inside the liposomes is flowed over the sensor to ameliorate artifacts due to buffer perfusion. Second, an “activating” buffer is flowed over the sensor to initiate transport. Capacitive coupling between the liposomal membrane and the surface-supported membrane on the electrode allows for measurement of ion transport currents (“on-currents”). Transport proceeds until a steady state is obtained where sufficient membrane potential opposes further net transport. Third, the non-activating buffer is reapplied, and the reverse transport process returns the sensor to its initial state. During this phase, “off-current” transport proceeds in the opposite direction of the on-current, driven by both chemical and electrical gradients.

Fig. 1 shows representative traces from our assay for all three stages of the experiment. First, non-activating buffer equilibrates the liposomes with a known concentration of proton and guanidinium. In the second stage, the activating buffer sets a 2-fold proton gradient to initiate transport. In Fig. 1A, there is no guanidinium gradient, so the outward-facing proton gradient should drive guanidinium into the liposomes. The negative on-current indicates that charge is moving out of the liposome, consistent with 2H^+^:1Gdm^+^ antiport. In Fig. 1B, the activating buffer sets an 8-fold outward-facing guanidinium gradient in addition to the 2-fold outward-facing proton gradient. Under these conditions, the large guanidinium gradient drives uphill proton transport into the liposomes, resulting in a positive on-current. In the third and final stage, current flux in the opposite direction from the on-currents is observed when non-activating buffer is re-applied to the sensor, as expected for reversible transport. While the off-currents in our experiments behaved as expected qualitatively, it is impossible to accurately quantify the electrochemical gradients during this third stage of the experiment. We therefore use only the on-currents for analysis.

**Fig. 1.**
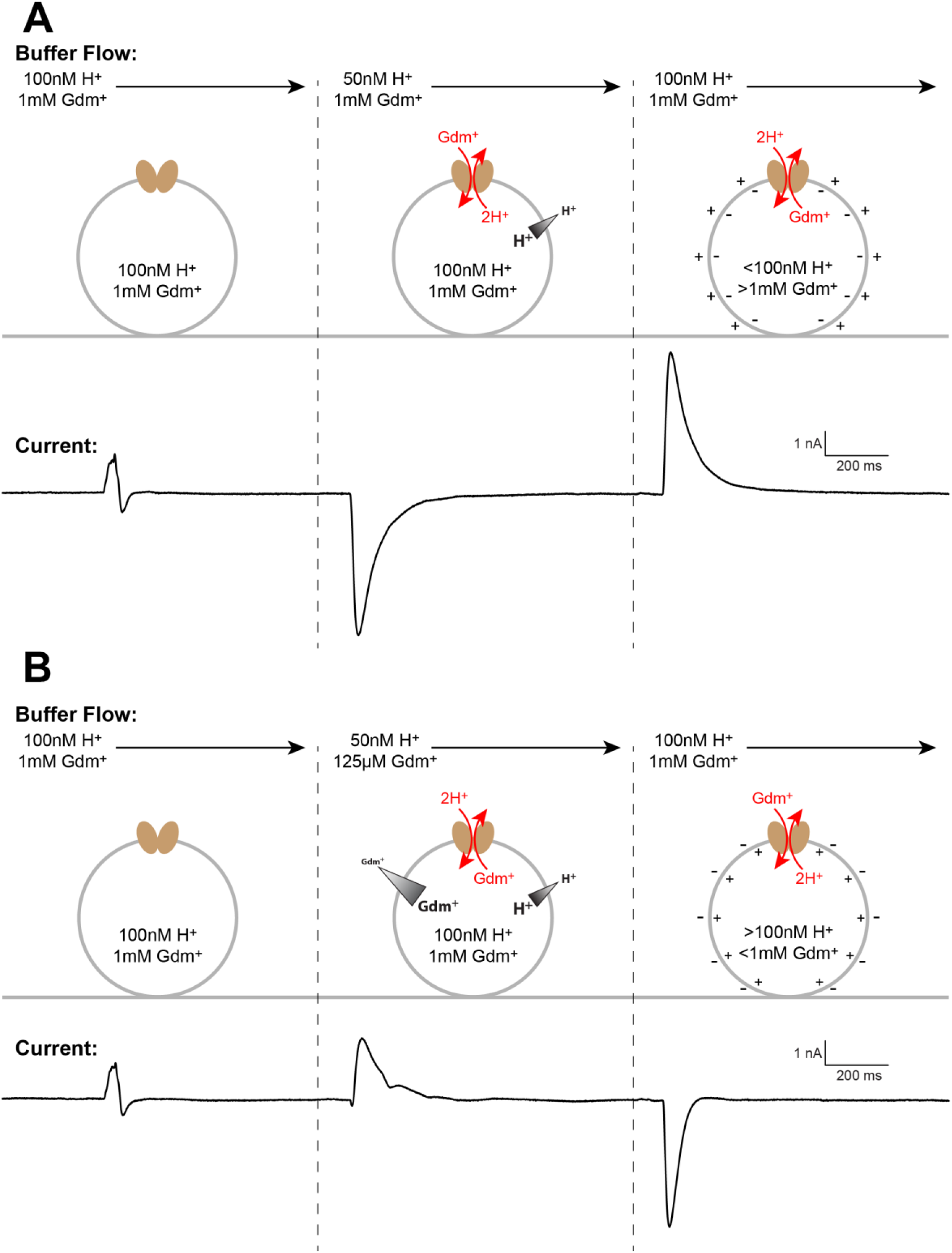
Assay scheme. Each assay consists of three stages of buffer perfusion. In the first stage, the “non-activating” buffer is identical to the internal buffer of the liposomes. In the second stage, the “activating” buffer sets a proton gradient and the guanidinium gradient is varied, leading to the transport on-current. In the third stage, “non-activating” buffer is reintroduced, and the transport off-current is observed as the liposomes return to their initial state. For this assay, only the on-current is quantified, so the remainder of the figures only display the second stage of buffer perfusion. **(A)** The activating buffer sets a two-fold proton gradient but no guanidinium gradient. The proton gradient drives guanidinium transport into the liposomes in exchange for two protons. Net charge is transported out of the liposomes, creating a negative current. **(B)** The activating buffer sets a two-fold proton gradient and an eight-fold guanidinium gradient. The guanidinium gradient drives uphill proton transport into the liposomes, creating a positive current.

For all of our assay conditions, we analyzed the results using two methods. SSME assays commonly quantify the peak on-current, a parameter that is related to the turnover rate of the transporter (Bazzone et al., 2017a; Garcia-Celma et al., 2010; Kelety et al., 2006; Schulz et al., 2008; Zuber et al., 2005). This parameter should reflect the direction and magnitude of transport that occurs as a result of the applied concentration gradients, and analysis of peak current as function of the chemical potential ratio (Fig. 2) should yield the reversal potential. However, peak current is ultimately a kinetic parameter, while transport stoichiometry is a thermodynamic quantity. We therefore considered a second method of determining reversal potential by analyzing the integrated current, representing the total net charge movement, as a function of the chemical potential ratio (Fig. 3). Before proceeding to this analysis, we performed several critical controls.

**Fig. 2.**
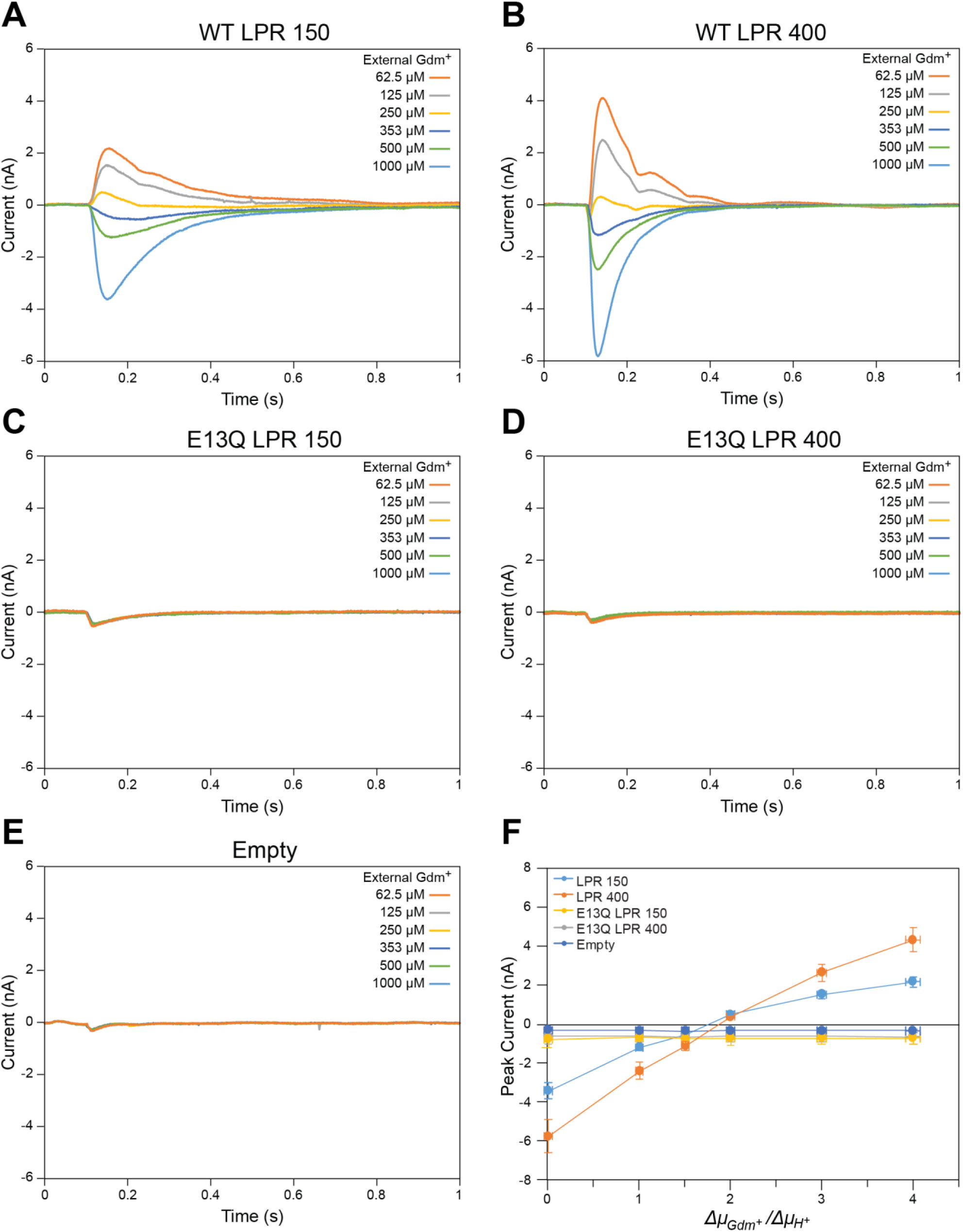
Peak current analysis is ambiguous. Representative current traces for each external guanidinium concentration are shown for WT-Gdx proteoliposomes with an LPR of 150 **(A)** and an LPR of 400 **(B)**, E13Q-Gdx proteoliposomes with an LPR of 150 **(C)** and an LPR of 400 **(D)**, and empty liposomes **(E). (F)** Plotting peak current against the chemical potential ratios yields a null current at a stoichiometry between 1.5 and 2H^+^/Gdm^+^. Data points represent average values obtained from four sensors for each (proteo)liposome sample. Error bars represent standard error of the mean.

**Fig. 3.**
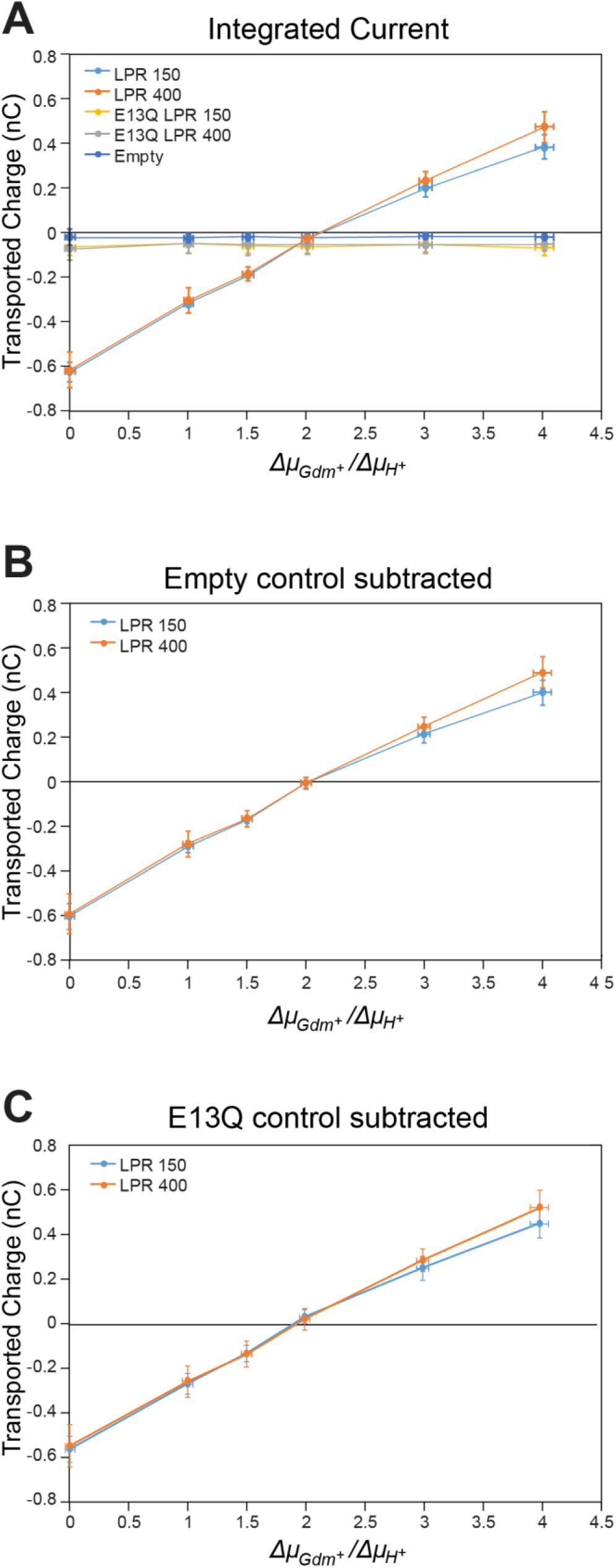
Integrated current analysis yields the correct stoichiometry. **(A)** Plots of integrated currents vs potential ratios for both LPRs of WT-Gdx, both LPRs of the transport-dead mutant E13Q-Gdx, and empty liposomes. Internal pH was 7.00 and internal guanidinium concentration was 1 mM. Null transport occurs at the published 2H^+^/Gdm^+^ stoichiometry regardless of whether the empty liposome integrated current is subtracted **(B)**, the dead-mutant integrated current is subtracted **(C)**, or simply the raw data from the WT integrated current is analyzed **(A)**. Data points represent average values obtained from four sensors for each (proteo)liposome sample. Error bars represent standard error of the mean.

First, we performed two experiments to ensure that the observed signal was not due to solution exchange and to assess the magnitude of solution exchange artifacts. We performed the assay using sensors prepared with both proteoliposomes reconstituted with the non-functional mutant E13Q-Gdx (Fig. 2C,D) and liposomes that were prepared with an identical reconstitution process but without protein (Fig. 2E). Solution exchange artifacts for both negative controls were of similar magnitude and small relative to transport signals for WT-Gdx (Fig. 2A,B). As an additional control, we prepared sensors from liposomes with different lipid to protein ratios (LPR) but with the same total amount of lipid (Fig. 2 A,C and B,D). Altering the protein concentration in this manner will not alter the thermodynamics of transport but will affect transport kinetics as well as pre-steady state currents related to electrogenic partial reactions, such as substrate binding or protein gating. Since we are interested in transport stoichiometry, it is important to check that the currents reflect the full transport cycle and not a partial reaction. Integrated currents for WT-Gdx are independent of LPR (Fig. 3), as expected if the currents correspond to transport.

Having confirmed that the currents reflect transport with the appropriate controls, we then compared the reversal potential using both peak current and integrated current analysis. The peak current reverses near the expected potential but plotting the peak current as a function of chemical potential ratios cannot unambiguously assign the stoichiometry (Fig. 2F). However, plotting the integrated on-current as a function of chemical potential ratio yields the published 2H^+^:1Gdm^+^ transport stoichiometry (Fig. 3A). This result is unchanged when either the E13Q (Fig. 3B) or empty liposome (Fig. 3C) controls are subtracted. This confirms that the thermodynamic parameter of integrated current is the better metric for determining transport stoichiometry in this assay.

Finally, one of the advantages of SSME over traditional liposomal assays is that a single SSM sensor is capable of dozens of measurements, drastically reducing sample requirements. Furthermore, it is well established that a single sensor can be used to screen different (external) activating buffers over several experiments (Bazzone et al., 2017a; Schulz et al., 2009). We wished to test whether it was possible to reliably change the *internal* buffer solution of the liposomes adsorbed to the sensors, which would expand the number of conditions that could be tested on a single sensor, further increasing the throughput of this method. We prepared sensors at pH 7.00 with 1 mM guanidinium in the same manner as in the experiments in Figs. 1-3. We then performed a series of rinses of pH 6.70 1 mM guanidinium buffer while monitoring the current using the instrument. After about 3mL total of rinse volume, the currents stabilized near zero (Fig. 4A). We proceeded to perform the reversal potential assay with a nominal internal pH of 6.70 and activating buffers at pH 7.00. The plot of integrated current against chemical potential once again yielded a transport stoichiometry of 2H^+^/Gdm^+^ (Fig. 4C), confirming that the internal pH was 6.70, as intended. This demonstrates that a single sensor can be used to test multiple internal substrate conditions. Altogether, this assay allows rapid data collection under the many different conditions required for transport stoichiometry determination while using a minimal amount of material.

**Fig. 4.**
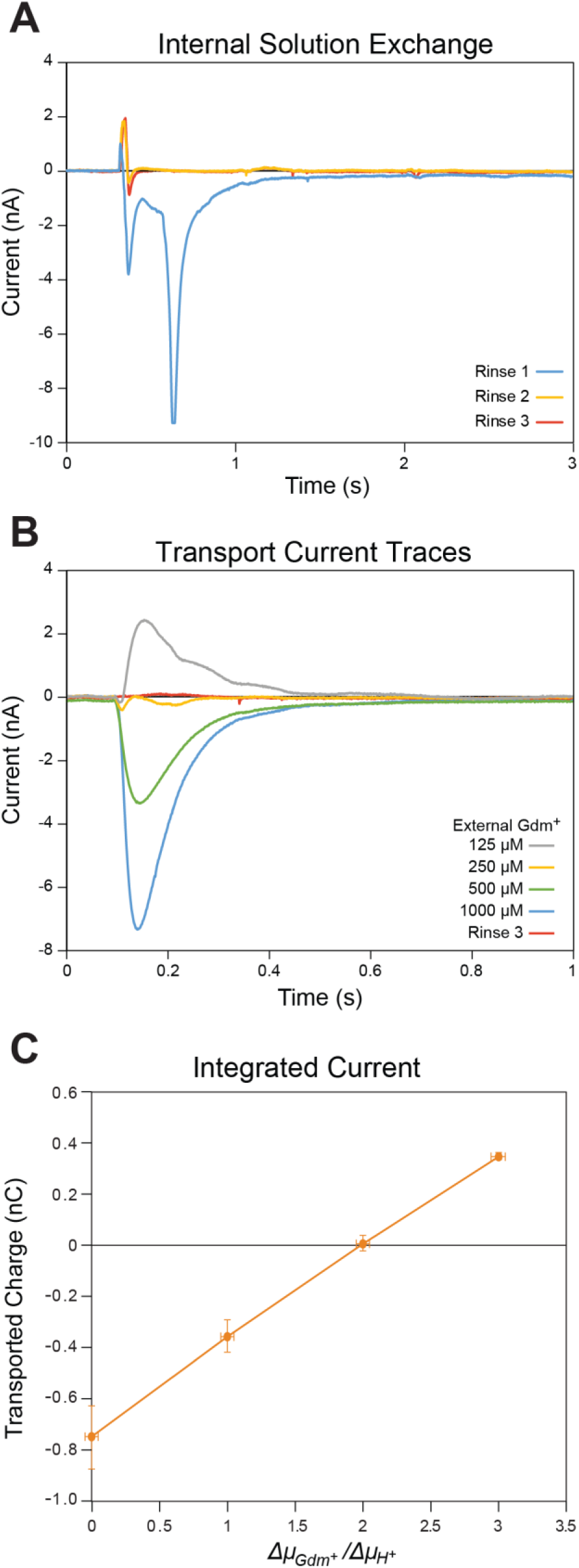
Internal liposome contents can be changed on the sensor. **(A)** Plots of currents of successive rinses to change the internal buffer to pH 6.70 and 1 mM guanidinium. **(B)** Representative current traces for each external guanidinium concentration with an LPR of 400. **(C)** Plots of integrated currents vs potential ratios for an internal pH of 6.70 and internal guanidinium at 1 mM. The LPR is 400 and integrated current of E13Q liposomes were subtracted. Data points represent average values obtained from four sensors for each sample condition. Error bars represent standard error of the mean.

## Discussion

Transport stoichiometry is a key determinant of transporter function, and accurate determination of transport stoichiometry can provide crucial insight into transporter mechanism. The current methods for measuring transport stoichiometry are time consuming and technically difficult, and as a consequence, transport stoichiometry has been quantitatively measured for only a small fraction of transporters that have been functionally or structurally characterized. Many of the transporters that have had their transport stoichiometry characterized have only been tested at a single experimental condition, which can lead to an assumption of a single transport stoichiometry that may not always be true (Grabe et al., 2020; Nguitragool and Miller, 2006). Our SSME assay removes many of the technical obstacles for stoichiometry determination and will facilitate measurement of transport stoichiometry for more transporters under a broader array of experimental conditions.

Apart from improvements to signal detection and throughput, the SSME assay may also allow for more reliable determination of transporter stoichiometry. In traditional liposomal assays, the internal buffer conditions must be set during reconstitution and cannot be checked for accuracy without expending some of the proteoliposomal sample in control reactions, further increasing the time and amount of sample required for the assay. In contrast, the SSME setup allows for repeated measurements on a single sample. All that is required to ensure that internal concentrations are accurate is to measure the current as “non-activating” (internal) buffer flows over the sensor. If the internal ion and substrate concentrations differ from the intended concentrations, a transport current will be evident, but no current will occur if the internal concentrations are matched to the known external buffer (Fig. 4A). This property can further be exploited to change the internal concentrations of the liposomes and test a wider variety of experimental conditions on the same sample.

As with any method, the reliability of this assay requires proper controls and assay conditions. SSME is sensitive enough to detect charge displacement due to conformational changes of proteins in the membrane (Bazzone et al., 2016), ion-transporter binding events (Călinescu et al., 2016, 2014), and solution exchange artifacts (Bazzone et al., 2017a), so it is important to distinguish transport currents from these other potential sources of current through proper controls. Negative controls should be used to estimate background current due to solution exchange artifacts. For Gdx, it is possible to abolish transport activity through a single mutation (E13Q), but transport-dead mutants are not available for every transporter. In such cases, “empty” liposomes which have undergone a simulated reconstitution process without protein may be used instead. The simulated reconstitution process is crucial, as the addition and subsequent removal of detergent can greatly affect the integrity of the lipid bilayer. As an additional control, sensors should be prepared with different LPRs, changing the protein concentration but keeping the lipid concentration constant. Integrated currents due to binding or conformational changes will change with protein concentration, but integrated transport currents will not, as they depend only on the transport potentials set by buffer exchange. In our results, there was no significant difference between the integrated currents for sensors prepared from LPRs of 150 and 400. However, it is interesting to note that at large substrate gradients, the integrated currents begin to diverge, with less transport observed for the LPR 150 sensors versus the LPR 400 sensors. Increasing the concentration of protein in the membrane (lowering the LPR) should increase the maximum rate of transport that can be achieved by a sensor, but it may also increase defects in the membrane, increasing membrane permeability and hindering maintenance of ion or substrate gradients. This suggests that increasing the LPR may increase the accuracy of the results in this assay, so long as sufficient signal to noise is maintained.

As our appreciation for the complexity of transporter mechanisms grows (Bazzone et al., 2017b; Bozzi et al., 2019; Parker et al., 2014; Robinson et al., 2017), it is increasingly important to be able to measure transporter stoichiometry. The SSME assay reported here addresses several key limitations of traditional reversal potential assays for measuring the stoichiometry of transporters. By directly measuring transported charge, this assay’s detection method is applicable to any electrogenic transporter that can be functionally reconstituted into liposomes. By allowing repeated measurements of the same sample, this assay reduces sample requirements and vastly increases throughput. Taken together, these improvements will allow for more routine measurements of transporter stoichiometry, facilitating in-depth characterization of transporter mechanism and function.

## Materials and Methods

### Sample Preparation

To minimize solution exchange artifacts, the buffers used for size exclusion chromatography, reconstitution, and electrophysiology steps had the same salt composition: 50 mM MES, 50 mM MOPS, 50 mM bicine, 100 mM NaCl, and 2 mM MgCl_2_. Buffer pH values were carefully adjusted using only NaOH to ensure that internal and external Cl^−^ concentrations were identical for all measurements.

WT- and E13Q-Gdx were expressed in *E. coli* from a pET15b vector and purified as previously described for the homolog EmrE (Morrison et al., 2011). Briefly, cells were lysed after overnight induction, and the protein was solubilized in 40 mM decylmaltoside (DM). Solubilized protein was run over a Ni^2+^-affinity column and eluted in 400 mM imidazole. The N-terminal hexahistidine tag was removed by overnight thrombin cleavage, and cleaved protein was further purified using size-exclusion chromatography.

WT-Gdx or E13Q-Gdx was reconstituted into POPC proteoliposomes at a lipid to protein ratio (LPR) of either 150:1 or 400:1 Gdx monomer (300:1 or 800:1 per functional dimer) in a pH 7.0 buffer. As an additional negative control, POPC liposomes were put through a simulated reconstitution process without protein. Detergent was removed with Amberlite XAD-2. Reconstituted liposomes were aliquoted and flash frozen. Immediately prior to measurements, liposome samples were thawed, diluted 2-fold in pH 7.00 buffer containing 2 mM guanidinium, and briefly sonicated. 10 μL of liposomes at a lipid concentration of 1.4 μg/μL were then used to prepare sensors for each sample condition.

### SSME Data Acquisition and Analysis

All electrophysiology measurements were recorded on a Surfe2r N1 solid-supported membrane-based electrophysiology instrument from Nanion Technologies GmbH (Munich, Germany). Prior to recording any transport measurements, sensors were rinsed with at least 1 mL of non-activating (internal) buffer while recording currents to ensure a flat baseline. Transport recordings occurred in three one-second stages according to Figure 1 with 200 μL/s buffer perfusion. After each transport measurement, sensors were again rinsed with 1 mL of non-activating buffer.

Initial data analysis was performed using the Surfe2r N1 instrument-specific analysis software from Nanion. The final 200 ms of non-activating buffer perfusion was averaged to obtain the baseline. Both peak current and integrated current data was obtained solely from the activating buffer perfusion. For transport currents with both positive and negative components (e.g., the 250 μM trace in Fig. 2A,B), the peak with the largest absolute value was recorded as the peak current. The entire stage of activating perfusion was integrated to obtain the integrated current values.

Four sensors were prepared for each sample (WT LPR150, WT LPR400, E13Q LPR150, E13Q LPR400, and empty liposomes). Reported data points consist of the average peak or integrated current of these four technical replicates. Transport behavior was highly consistent between sensors, so no additional replicates were performed. Error bars represent the standard error of the mean. No outliers were observed, and no data was discarded.

### Derivation of transport equations

Assume that transport proceeds according to the stoichiometric transport reaction:

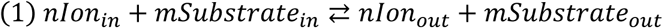

For symport (co-transport) *n* and *m* are the same sign, while for antiport (exchange), they are opposite signs. The chemical potential of ion and substrate across the membrane is given by:

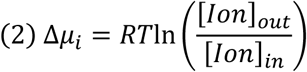

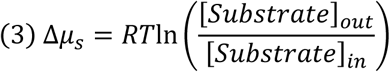

In the absence of a membrane voltage, the free energy for the coupled transport reaction is given by:

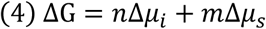

When ΔG = 0,

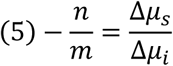

Thus, plotting transported charge against *Δμ*_*s*_*/Δμ*_*i*_ gives an x-intercept at -*n/m*, allowing determination of the transport stoichiometry.

## Acknowledgements

We would like to thank Dr. Maria Barthmes and Dr. Andre Bazzone for their support of the assay development. This work was supported by the National Institutes of Health Grant 1R01GM095839 (KHW) and a NIH National Research Service Award T32 GM007215 (NET). The authors declare no competing financial interests.

